# Peppy: A Virtual Reality Environment for Exploring the Principles of Polypeptide Structure

**DOI:** 10.1101/723155

**Authors:** David G Doak, Gareth S Denyer, Juliet A Gerrard, Joel P Mackay, Jane R Allison

## Abstract

A key learning outcome for undergraduate biochemistry classes is a thorough understanding of the principles of protein structure. Traditional approaches to teaching this material, which include two-dimensional (2D) images on paper, physical molecular modelling kits, and projections of 3D structures into 2D, are unable to fully capture the dynamic, 3D nature of proteins. We have built a virtual reality application, Peppy, aimed at facilitating teaching of the principles of protein secondary structure. Rather than attempt to model molecules with the same fidelity to the underlying physical chemistry as existing, research-oriented molecular modelling approaches, we took the more straightforward approach of harnessing the Unity video game physics engine. Indeed, the simplicity and limitations of our model are a strength in a teaching context, provoking questions and thus deeper understanding. Peppy allows exploration of the relative effects of hydrogen bonding (and electrostatic interactions more generally), backbone ϕ/ψ angles, basic chemical structure and steric effects on polypeptide structure in an accessible format that is novel, dynamic and fun to use. As well as describing the implementation and use of Peppy, we discuss the outcomes of deploying Peppy in undergraduate biochemistry courses.

**STATEMENT:** Protein structure is inherently dynamic and three-dimensional, but traditional teaching tools are static and/or two-dimensional. We have developed a virtual reality teaching tool, Peppy, that facilitates undergraduate teaching of the principles of protein structure. We outline how Peppy works in terms of how it is used and what goes on ‘under the hood’. We then illustrate its use in undergraduate teaching, where its playful nature stimulated exploration and, thus, deeper understanding.

## Introduction

The principles of protein structure are a threshold learning outcome for fundamental undergraduate biochemistry courses. Understanding the structures and conformational preferences of amino acids, and their capacity to make non-bonded interactions such as hydrogen bonds and ion pairs is fundamental to appreciating the formation of regular secondary structure elements such as α-helices and β-sheets. Functional competence with protein structure requires not only committing these principles to memory, but also gaining a sense of how a protein’s three-dimensional (3D) structure emerges from the interplay of the underlying physical and chemical characteristics.

Traditional approaches to teaching students about protein structure include textbook 2D images on paper through to physical molecular modelling kits, and to projections of 3D structures into 2D such as stereograms and protein molecular graphics programs. Although these are all useful in different ways, all suffer from limitations. Proteins are inherently 3D objects, and thus any 2D representation will fail to provide a complete picture. Physical models are 3D, but are fragile and time-consuming to assemble and disassemble. Moreover, the behaviour of proteins is intrinsically dynamic and results from the interplay of a host of physicochemical forces, which are difficult or impossible to represent in rigid models or on paper. 3D molecular visualisation software packages are typically only used in senior classes, and even these tools are generally limited to observing pre-determined, static structures and still require the mind to derive a 3D understanding from a 2D computer screen.

For these reasons, an alternative approach that might assist students in understanding the underlying principles is to use a virtual reality (VR) environment. VR is intrinsically 3D and allows both representation of dynamic behaviour and ‘hands-on’ manipulation by the user.

The potential benefits of teaching protein structure using an interactive VR approach have already been reported (23; 11; 5; 2). The particular strengths of VR are not limited to its novelty or connection to gaming, but are derived from the physical involvement and visual immersion of the user, which is facilitated by having a head-mounted display and the availability of six degrees of freedom (6DoF; translation along and rotation about each of three orthogonal axes).

There are many existing examples of the use of VR technology to visualise experimental data (8; 21) and facilitate the investigation of cellular(8) (e.g. http://thebodyvr.com) and molecular (22; 12; 6; 18; 1; 4; 24; 5; 13; 15; 9; 14) (also e.g. http://nanome.ai, https://gwydion.co, https://research.nanosimbox.io) scenarios. Most, however, are targeted at researchers rather than undergraduate students and tend to deliver material rather than encourage its production. While this does not prevent their use in teaching, the design goals of a tool aimed at teaching are quite different to those of a tool aimed at researchers. Effective teaching requires a fun and intuitive environment that encourages self-directed and creative engagement and leads the students to ask questions; thus, a degree of fallibility is desirable and genuine exploration and productive failure is essential.

Although the case for using VR in teaching molecular processes is compelling, the impetus to create specialised applications is somewhat reduced by the fact that deployment of VR to large undergraduate classes is limited by a lack of specialised, high-throughput facilities. Furthermore, even when suitable software exists, deployment requires agility in course management to allow rapid introduction of into the curriculum.

We present here a VR tool, ‘Peppy’, aimed at facilitating the teaching of the principles of protein structure to undergraduate classes. Peppy allows exploration of the relative effects of hydrogen bonding (and electrostatic interactions more generally), backbone ϕ/ψ angles, basic chemical structure and steric effects on the resulting polypeptide structure. Additionally, we describe the prototyping of Peppy in undergraduate biochemistry courses at the University of Sydney, which possesses a dedicated VR facility, the Immersive Learning Laboratory (ImLL). This, along with careful yet adventurous course design and management, overcame the aforementioned issues with deploying VR in teaching.

## Methods

### Development Strategy

Our goal, and thus our design approach, was to model a traditional physical ball and stick representation of molecular (peptide) structure with a dynamism that would enhance student engagement but with a realism that would ensure quality learning. To achieve this, we took advantage of the existence of video game development engines that have at their core a robust physics engine and 3D rendering, whilst also offering the ability to rapidly prototype an application, thus allowing rapid and agile cycles of design and testing.

Achievement of our goals does not require the same degree of realism as the force fields used in molecular dynamics simulations, nor does the visual representation need to be as sophisticated as existing molecular visualisation such as Pymol (19) and VMD (7). Indeed, in contrast to the latter tools, the refresh rates and rendering required for a pleasant user experience impose a further limitation on functionality (10). Peppy does not, therefore, represent a robust, fully-featured molecular dynamics simulation, but rather, the simplest possible functional model of a polypeptide chain within a game engine. However, the fidelity of the physics within the game engine is very high, and the underlying computational methods are not dissimilar to those used in molecular dynamics simulations. Crucially, the end result is dynamic with an intuitive game-like interface that is highly interactive in real time.

### Implementation

Peppy was created using the Unity game engine (https://unity3d.com). We note that the units are those used in Unity and are in general at a human scale, e.g. distances are in metres, weights are in kilograms, as is standard practice in game development. Geometry and prefabricated (prefab) components were created within the Unity editor and associated code is written in C#. Some code components are licensed from the Oculus Software Development Kit (SDK). The source files and compiled executables for Peppy are available at https://github.com/ddoak/peppy.

Peppy runs on any VR-capable desktop machine with Oculus Rift headsets and touch controllers, and is also available for Oculus Quest. To broaden its accessibility, it may also be run in a non-VR ‘flat screen’ mode without Oculus hardware. In this mode the user’s movement and interaction is controlled using mouse and keyboard.

### Sterics, geometry and rendering

Peppy describes the polypeptide chain at an all-atom level of detail, in keeping with standard molecular dynamics force fields. This representation is functionally implemented using Prefab GameObjects within Unity. A *prefab* is a user-defined reusable template comprising a hierarchical collection of components such as transforms, mesh renderers, rigidbodies, colliders and C# scripts, which define bespoke behaviours and properties. *Transforms* define the position, rotation and scale of an object; *mesh renderers* render the object in 3D at the position defined by the transform; *rigidbodies* are internally rigid objects that behave according to the laws of physics; and *colliders* define the shape of an object for the purposes of physical collisions. *Configurable joints* connect the rigidbodies. These are oriented such that the x-axis of the joint aligns with the bond and are locked in y and z so that they can only rotate around the x-axis.

Atoms, for instance, have individual spherical meshes for rendering in addition to fixed-radius hard spherical colliders that prevent interpenetration. The collider radii are derived from standard van der Waals radii (3) and are not adjusted for different chemical environments (**Supporting Information Table S1**). Attractive van der Waals forces and the effects of solvent are not modelled for simplicity.

A typical polypeptide fragment prefab in Peppy is effectively a united atom representation comprising a single rigidbody component with appropriate transforms and mesh geometry representing the associated atoms, fixed internal bond lengths and angles. Rigidbodies (and hence the prefab units built from them) are connected by configurable joints between anchor points coincident with the appropriate bonded atom centres, which have only one permitted DoF (axial rotation).

The colliders of bonded atoms are permitted to intersect. However, within a prefab unit, these colliders combine to form compound colliders that do not self-interact. For adjacent prefab units, connected by a configurable joint, collider interactions are explicitly turned off. Thus, effectively, in keeping with the exclusions common to molecular dynamics force fields, the van der Waals interactions between bonded atoms are excluded.

The colliders can be switched on and off by the user to allow exploration of the restrictions on conformation imposed by steric hindrance.

Bond lengths and angles are encapsulated by either the fixed internal geometry of the prefab transforms (**Supporting Information Table S2**) or the parameters for the configurable joints (**Supporting Information Tables S3**, **S4**). Within the sidechains, departures from idealised sp^2^ (trigonal) geometry are specified explicitly (**Supporting Information Table S5**).

The radii of the visible atomic render mesh spheres are scalable, which allows the user to transition smoothly between ball and stick and Corey-Pauling-Kulton (CPK) shell representations. We note however that the collider radii do not change, only the rendering.

Rendered bonds (grey cylinders) are entirely cosmetic – they are simply a fixed cylindrical mesh geometry connecting atoms. Bonds joining rigidbody units (see below) are aligned and thus generally coincident with the main axes of the corresponding configurable joint. If the configurable joints are highly strained (e.g. if the user pulls the polypeptide backbone apart), however, there may be a noticeable mismatch between the rendering and the underlying physics.

The masses of all prefab unit rigidbodies are scaled appropriately to represent the combined mass of their constituent atoms (**Supporting Information Table S1**).

### Backbone architecture

The polypeptide backbone is built from three types of prefab unit – N-H, H-Cα-R, and C=O – with fixed internal bond lengths and angles (**Figure 6**). The configurable joints connecting the backbone prefab units represent the polypeptide backbone covalent bonds. They are fixed for the peptide bond but free to rotate for the central bond of the ϕ and ψ dihedral angles, providing the minimum required rotational DoF (two rotations per residue).

Each configurable joint has a target dihedral angle value and an associated spring force (torque). Both can be controlled by the user. Target dihedral angle values can be chosen from a Ramachandran plot for all or selected amino acids. If the torque is non-zero, the dihedral is driven toward the target value. The torque values are not representative of real intra-molecular forces; rather, they allow the user to manipulate the polypeptide backbone toward particular conformations. The scaling for the torque is empirical and was tuned during development to give a range of values that allow the user to explore secondary structure and steric hindrance.

### Sidechain architecture

Amino acid residues are by default created with single-atom dummy ‘R’ side chains but may be selectively mutated at runtime to any of the standard twenty amino acid sidechains. Inter-residue disulfide bonds may also be created by selecting pairs of cysteine residues to join.

Specifying a particular amino acid sidechain replaces the generic R (magenta sphere) of the backbone (H-Cα-R) with appropriate connected prefab units (*e.g.*, CH_2_, NH_3_^+^, OH, etc.). As for the polypeptide backbone, groups of atoms are unified for simplicity, with the divisions generally located on rotatable bonds, and each atom is described by an atom-specific collider. Key parameters describing the side chain architecture are provided in **Supporting Information Table S5**.

### Hydrogen bonds

Hydrogen bonds between backbone donor atoms (H-N) and acceptor atoms (O=C) are modelled explicitly so that they can be tuned independently of other electrostatic interactions, thus enabling students to explore their effect on protein secondary structure. Candidate hydrogen bonds are identified using a ‘spherecast’ test, which sweeps a cylinder away from the donor atom, extending along the direction of the amide bond, to search for acceptor atoms (O=C). This test is interrupted by the presence of other atoms to prevent tunnelling. The radius (0.05 m) and length (0.3 m) of the test cylinder were empirically tuned to facilitate generation of the predicted hydrogen bonds in regular secondary structure elements.

If the test locates a candidate acceptor, the relative distance between the donor and acceptor is switched linearly from 1 if the acceptor is located as being at a distance of 0 m from the donor atom to 0 if the acceptor is located at the furthest end of the cylinder (0.3 m).

If the test locates a candidate acceptor, a hydrogen bond is modelled with three spring joints (20) (**Figure 7**). The joints are modelled with a flat-bottomed, damped harmonic potential:

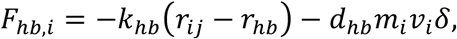

where *r*_*ij*_ is the current distance between the two atoms, *i* and *j*, that the spring joint connects, *r*_*hb*_ is the ideal length of the spring, *k*_*hb*_ is the hydrogen bond spring force constant, *d*_*hb*_ is the hydrogen bond spring damping constant, *m*_*i*_ is the mass of atom *i*, *v*_*i*_ is the instantaneous velocity of atom *i*, and the Kronecker delta, *δ*, is defined as:

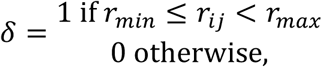

where *r*_*min*_ and *r*_*max*_ are the minimum and maximum of a distance range within which the spring does not act. The values of the ideal spring lengths, force constants and damping constants are provided in the **Supporting Information Table S6**. Using three springs encourages linearity of the four atoms involved (N – H … O – C). Essentially, the springs attract the hydrogen atom to the oxygen atom whilst repelling the nitrogen atom from the carbon atom. Because the lone pair geometry of the acceptor oxygen atom is not modelled, the overall effect is to favour a linear hydrogen bond. Intra-chain hydrogen bonds between residues *i* and *i* ± *n*, where *n* ≤ 2, are explicitly excluded.

The springs obey Hooke’s law, with the force proportional to the displacement from the target length. This would result in extremely large forces at large inter-atomic distances, which would be unrealistic due to the 1/*r* distance dependence of the electrostatic forces that underlie hydrogen bonding and solvent screening effects, but do have the advantage of being robust to user interaction. The simple linear switching function described above, which makes the spring proportionally weaker as the donor and acceptor atoms/groups move further apart, was therefore introduced to stop the spring forces from becoming too large relative to the other forces in the model when they are stretched.

Active hydrogen bonds are visualised through a simple animated particle effect that is continuously updated to be oriented directly from the H-N donor toward the O=C acceptor.

The scale for the spring forces is completely empirical and can be adjusted by the user. The lower end of the scale represents no hydrogen bond formation, while the upper end of this scale represents unrealistically strong hydrogen bonds. This deliberate choice allows the user to experiment with manipulating robust secondary structure elements.

A clear limitation of this current approach is that each hydrogen bond is independent of the presence of other hydrogen bonds. Consequently, multiple donors can hydrogen bond to a single acceptor. Although this is observed in experimental structures and in more sophisticated molecular dynamics simulations, it is overly prevalent in the current version of Peppy. The independence of hydrogen bonds does not prevent the occurrence of higher-level cooperativity of hydrogen bond formation, for instance, in the “zipping” together of β-strands into a β-sheet.

### Electrostatic interactions

The electrostatic interactions between a subset of polar atoms are modelled explicitly via a simplified Coulombic potential (**Figure 8**). The number of atoms with partial charges is restricted in comparison to more accurate molecular dynamics simulations for performance reasons as outlined below. Partial charges, *q*, are assigned to backbone amide hydrogen atoms, carbonyl oxygen atoms, and the sidechain atoms of amino acids that would be ionised at physiological pH (arginine lysine, glutamate, aspartate, histidine). The values of the partial charges (**Supporting Information Table S7**) are derived from the GROMOS 54A8 forcefield parameters (16; 17). The total electrostatic force on a partially charged atom is given by

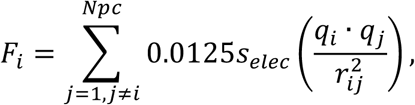

where *N*_*pc*_ is the total number of atoms with partial charges, 0.0125 is an empirical scaling factor, *r*_*ij*_ is the distance between atoms *i* and *j*, and *i* and *j* do not reside within the same prefab unit. *s*_*elec*_ is the electrostatic strength and can be adjusted by the user in the range [0,100] to investigate the contributions of electrostatic forces to peptide structure. The resultant force is applied to the parent rigidbody at the position of atom *i*.

Electrostatic interactions are visualised through animated particle effects. Each charged unit has a coloured (red/blue) particle system that emits radially from a spherical volume around the atom. The number, size and acceleration of the particles is scaled according to the magnitude and direction of current resultant electrostatic force on the atom. This effect is deliberately theatrical and is intended to be arresting for teaching purposes.

In order to contain the computational cost, the electrostatic interactions are computed and resulting forces applied at lower frequency than the game physics (10 *vs* 90 iterations per second). Moreover, because the current implementation does not use neighbour lists or distance exclusions, the cost of calculating the electrostatic interactions increases rapidly (О*N*_*pc*_^2^) as the number of partially charged atoms increases.

### Direct interaction

Prefab atom groups can be ‘grabbed’ and manipulated directly using the motion controllers. Direct grabbing is the routine way that object manipulation is implemented in many VR games. The grabbed prefab unit is directly attached to the user’s hand (controller), and thus inherits the hand position and rotation (transform), giving the user intuitive 6DoF control.

Whilst direct interaction is highly intuitive and responsive, it has some inevitable issues. Grabbing and manipulating a prefab unit effectively takes control of its transform away from the underlying physics governing the peptide behaviour. Reconciling the grabbed object movements with those of the other connected prefab units that are controlled only by the underlying physics introduces effectively unlimited forces/torques. It is therefore possible to inadvertently distort bond lengths and angles. In addition, large rapid movements have the potential to generate ‘explosive’ oscillations as the fixed time step simulated physics struggles to reconcile large instantaneous changes. It should be noted that this can also be a problem in more sophisticated molecular dynamics simulations, but may be rectified with the development of more sophisticated controllers.

### Remote interaction

Prefab atom groups can also be interacted with at a distance. Pointing a motion controller at an atom group highlights the group which can then be ‘tractor beamed’ directly towards or away from the user. This functionality is further extended with a ‘remote grab’ interaction, which allows the user to intuitively push, pull and tangentially drag an atom group at a distance. This feature works by calculating an appropriate translational vector from the user’s motion controller gesture and applying an impulse force to the remotely grabbed object. Although direct grabbing is perhaps more intuitive for a new user, the remote-interaction approach turns out to be very valuable for manipulating polypeptides.

### Dynamics

The forces resulting from the underlying physics are always heavily damped. Without this, the prefab’s rigidbodies would be prone to acquiring large velocities particularly when being directly interacted with by the user. Damping is achieved by all rigidbodies having empirical drag factors enabled for translational and angular motions (**Supporting Information Table S8**). The overall scaling of these drag values can be adjusted by the user via a slider but can never be reduced to zero. Selected residues may be ‘frozen’, which sets the drag parameters for the associated prefab units to infinity.

‘Jiggle’ dynamics, notionally equivalent to thermal motion, are achieved by applying random impulse forces to each of the rigidbodies. The impulse forces are randomised every frame and the overall scale factor is empirical. These forces are applied to the centre of mass of the rigidbodies meaning that, as a consequence, prefab units (such as CH_3_ groups) do not spin as much as is observed in molecular dynamics simulations.

## Results

We first outline the principles and goals underlying our approach, and then describe the high-level functionality of and user interaction with Peppy, followed by details and outcomes of its deployment in undergraduate biochemistry classes. Although both our implementation and our testing to date are preliminary, we think Peppy shows great promise as a teaching tool. The code is publicly available *via* GitHub and we encourage others to try it out and provide us with feedback that we will harness to inform further development.

### Underlying principles and goals

Our goal was to create an environment that allowed students to engage with protein structure and dynamics and gain an understanding of how these are determined from the underlying physics and chemistry. Our fundamental philosophy was to encourage experiential learning about the conformational properties of polypeptides through play.

We have harnessed the gaming associations of VR, as well as its ability to provide 3D information at human scale, to enhance student engagement and ‘make learning fun’. To facilitate understanding and experimentation, absolute physical/chemical correctness is less important than usability. We therefore allow interactive alteration of just the major factors that influence secondary and tertiary structure. To maximise simplicity, the tunable factors are limited to those that have most impact on protein secondary and tertiary structure, namely residue types, ϕ/ψ angle values, hydrogen bonding and electrostatic interactions. These can be adjusted by the user between their fully off and fully on states; in the fully on state, that factor will dominate other factors that are not in a fully on state.

### High-level functionality

Peppy allows the user to create polypeptide chains that can then be ‘physically’ grabbed and manipulated in the virtual space. By pushing, pulling, twisting and ‘touching’ these molecules, higher order structures can be created or destroyed and their stability and properties investigated. The minimal game-like environment encourages self-directed creative engagement. Interaction is immediate and intuitive and is built on both the immersive nature of VR and the revolutionary interface possibilities afforded by fully tracked 6DoF motion controllers. Many of the low-level simulation parameters (*e.g.*, force constants, dynamics) are exposed to the user and can be manipulated directly and the consequential effects observed. Peppy is not intended to be a robust and detailed molecular dynamics simulation; however, it is highly effective as a representative sketch that allows the user to explore many of the emergent structural properties of proteins such as repeating secondary structural elements. Hydrogen bonds and electrostatic interactions are modelled in a simplified manner and are represented graphically by animated particle effects that visualise the dynamic forces involved.

Backbone (ϕ and ψ) dihedrals are visualised on an interactive Ramachandran plot and can be manipulated and monitored by the user. Amino-acid sequence can be easily altered in order to visualise and investigate the impact of sidechain conformations and steric properties. An in-game camera allows the user to record snapshots of their creations, in association with an avatar that projects their physical presence within the virtual environment.

It is also possible to run the application in a ‘flat’ non-VR mode – this presents the same innovative dynamic functionality but is limited by a more traditional mouse/keyboard interface. While losing the immersive nature of the VR environment, this mode allows students to explore the application on a standard Windows operating system without the requirement for VR hardware, thus broadening the penetration of the software.

### User interaction

Peppy includes a variety of methods for the user to interactively manipulate peptide molecules in real time. Most interactions can be controlled and reported though the main dashboard, which is shown in **Figure 1**.

**Figure 1.**
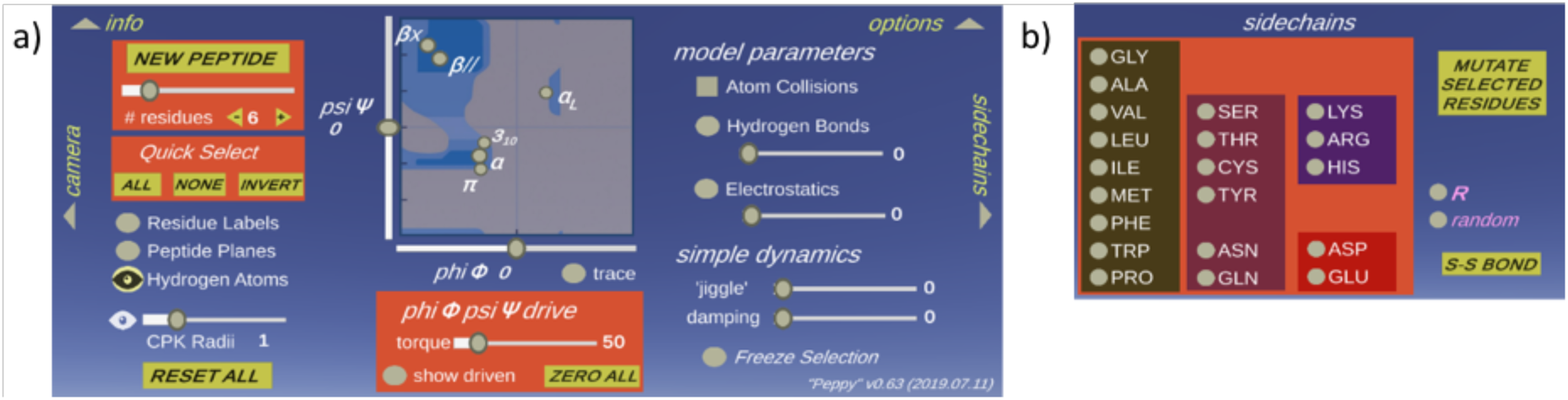
The Peppy interaction dashboard. (a) Main dashboard and (b) pop-out sidechain menu. Arrows near the edges can be ‘clicked’ to open out additional specialist menus. The *tunable parameters*, which are adjusted for selected residues using sliders, are hydrogen bond strength, electrostatic interaction strength, ϕ/ψ angle values, ϕ/ψ drive torque, visualisation of atomic radii, degree of ‘jiggle’ dynamics and damping of the dynamics. The *binary options*, which are turned on or off using radio buttons, are the calculation of forces due to atom collisions, the visibility of hydrogen bonds, electrostatic interactions, peptide planes, and hydrogen atoms, freezing selected residues, the illustration of the ϕ/ψ trace, highlighting of residues with driven ϕ/ψ, and numbering of residues in the Ramachandran plot. *Adjustable peptide properties* are the peptide size (number of amino acids) and the type of each amino acid type (via the pop-out *sidechain* menu).

In a standard interaction, a user first builds a peptide backbone of a particular length. They may then manipulate the torsional angles between the peptide units to create motifs of secondary structure, either though manual manipulation or by selecting areas on the Ramachandran plot. In addition, users may explore the influence of particular side-chain residues by altering (or “mutating”) particular R-groups.

#### Building a peptide

In first constructing a polypeptide unit, the user chooses the number of amino acids and a peptide with ‘dummy’ (R) side chains appears. These R-groups can subsequently be mutated individually or in groups to any amino acid type. There is also an option to randomly choose the amino acid type. Residue numbers are shown if the ‘Residue Labels’ radio button is selected. Visibility of peptide planes and hydrogen atoms can also be controlled in the same manner.

#### Selection

Individual residues can be selected by pointing a controller at a prefab unit and ‘clicking’ on any of its atoms. Contiguous prefab units can be selected by as a group, and all the residues in a peptide can be selected through buttons in the ‘Quick Select’ box. Once selected, the type of amino acid and the ϕ/ψ angles can be modified though the dashboard.

The peptide(s) and the menu can be moved closer or further away by remote interaction using the ‘tractor beam’. Peptides can be ‘grabbed’ directly through the user’s virtual hands, or ‘remote grabbed’ and directly pushed, pulled or tangentially dragged using tractor beam. The remote grabbing interaction is sufficiently intuitive and direct to allow the user to carry out sophisticated manipulations. Remarkably, the flexibility and smoothness of the molecular handling is sufficient to allow a peptide backbone to be tied in a knot using a single motion controller.

#### Physical effects

The contribution of steric effects can be switched on or off using the ‘Atom Collisions’ radio button. The effective size (radius) of each atom is fixed, but atoms can be visualised at different sizes by tuning atom size using the ‘CPK Radii’ slider.

The ϕ/ψ angle values are visualised on a 2D map modelled on the Ramachandran plot commonly used to analyse protein structure. Major secondary structure types are represented by selectable spots on the map. The ϕ/ψ angle values of a single or group of selected residues can be controlled by clicking on a position in the 2D map or by sliders on each axis of this map. The sliders direct the parameters of the configurable joints representing the ϕ/ψ bonds. The strength by which ϕ/ψ angle values are driven to the selected values is controlled using a slider which adjusts the apparent torque of the bonds.

The current ϕ/ψ angle values of each residue are dynamically displayed. Driven dihedrals can be visualised *via* a toggle switch, and the history of the ϕ/ψ angle values can be shown by selecting the ‘trace’ option.

#### Chemical properties

Atoms are coloured by type similar to commonly-used atom rendering standards, with carbon black, nitrogen blue, oxygen red, hydrogen white, sulfur yellow, and ‘dummy’ side chains magenta; the atomic radii depend on the atom type.

Although hydrogen bonds are a subset of electrostatic interactions, in Peppy they are modelled explicitly to allow students to explore their effect on protein secondary and tertiary structure independently of other types of electrostatic interactions. Both electrostatic interactions and hydrogen bonds can be visualised independently by checking their respective radio buttons. Similarly, the strength of both types of interaction can be scaled using a slider.

#### Dynamics

An essential part of Peppy is that the simulation is interactive and dynamically responsive in real time. By default, the peptide is static, but any forces added by the user (*e.g.* by grabbing and manipulating part or all of a peptide) cause a wave of movement through all the residues in response. The resultant forces are heavily damped to prevent anomalously large velocities occurring.

Random motion, in which each rigidbody experiences impulse forces of random direction that modify the inherited momentum, can be incorporated by switching on ‘jiggle’. This is notionally equivalent to thermal motion but no attempt is made to correlate the scale of the jiggle dynamics with a particular macroscopic temperature. The dynamics are also damped by scaling the translational and rotational motion. Both the size of the impulse forces and the degree of damping can be adjusted.

Selected residues may be ‘frozen’ to allow the user to easily lock conformations of regions the polypeptide to facilitate more directed manipulation elsewhere.

### Outcomes and Deployment

We deployed Peppy in undergraduate biochemistry classes at the University of Sydney in August 2018 and March 2019. The first cohort comprised a class of 80 biochemistry students (course code BCHM3082), most in their final semester at University. There was no assessment component; rather, we aimed to investigate user acceptance and capability, as well as the logistics of running large-class VR sessions in a dedicated facility. Interestingly, less than 5% of the cohort reported ever having worn a VR headset, but this did not appear to be an impediment to the majority of students. Encouraged by the reception, we carried out several iterations of development of Peppy before using it for a class of 85 3^rd^ year students in a course specifically related to protein structure and function (BCHM3072). As with the first cohort, these students were familiar with the fundamentals of protein secondary structure, and the majority had explored this in the 2^nd^ year foundational courses by constructing physical models of short polypeptides using Cochranes Orbit molecular building system (https://www.cochranes.co.uk/). Despite this, their depth of understanding for many of the concepts we consider necessary for fluency when discussing structural biology was largely still developing.

Students worked in pairs for about 90 minutes to complete a series of tasks using Peppy. The workflow was intended to take the students from the creation of short peptides, for the revision and exploration of basic peptide bond geometry, through to the construction of complex polypeptides, and the formation of hydrogen-bonded secondary structures (α-helices and β-sheets). Students were encouraged to investigate the effects of changing tunable-parameter values and amino acid side chains to see how this affected the secondary structure elements that they had made (**Figure 2**).

**Figure 2.**
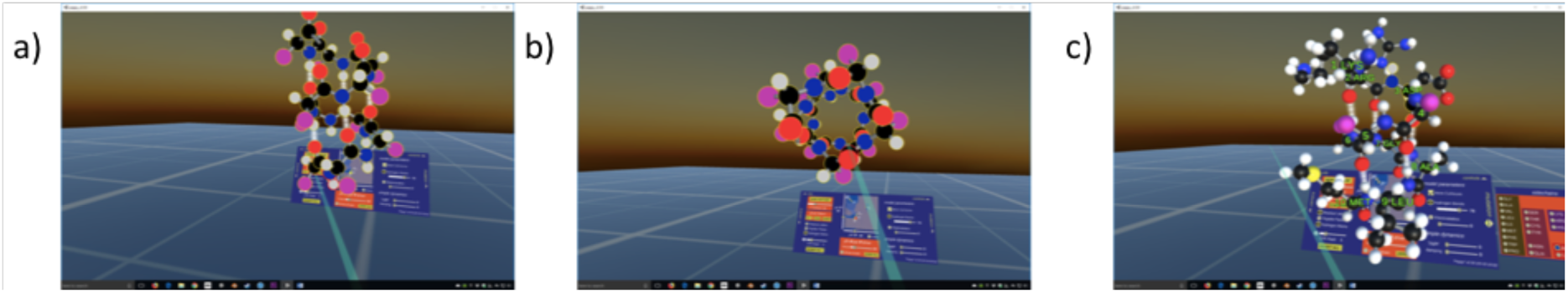
Examples of α-helices built by students. (a) angled view showing hydrogen bond formation in an α-helix built with dummy sidechains (magenta); (b) end-on view showing the positions of the side chains for the same α-helix; (c) an α-helix with post hoc specification of the amino acid side chains so that the uppermost residues are hydrophilic and the lower residues are hydrophobic. The students commented “*We initially tried to put extremely large residues in the helix like tryptophan*, *but we soon found out that this made the helix unwieldy. We therefore largely stuck to smaller residues like glycine*, *lysine and alanine. The R groups of the top section began to come close to ionically bond together as we put an aspartate and lysine/arginine group next to each other. We did this deliberately to see how rigid the helix was*, *and whether the ionic strength would pull apart the helix.*”

Anecdotally, we observed that working in pairs was exceptionally effective, with students taking turns to wear the headset or read and interpret the written instructions. This alternation of pilot and navigator fostered engagement, reduced the burden of wearing the headset for extended periods, and created a strong culture of inter-student support and desire to achieve mastery. It prompted not only discussion about which features of Peppy were exciting and which could be improved, but also led students to openly confess previous misconceptions and provide explanation to one another.

After completion of the workflow, students were encouraged to explore all the functions of Peppy and, almost unanimously, they enthusiastically built extravagantly complex polypeptides, experimented with the effects of different amino acid side-chains, and attempted to construct multi-chain tertiary structures such as β-barrels (**Figure 3**) and even real, small proteins (**Figure 4**). One student even attempted to recreate the active site of trypsin by arranging and orienting key resides in 3D space, taking directions from a representation of the structure in Pymol. As expected, this was far too ambitious but, as with all these cases, the struggle made the student appreciate the incredible complexity and beauty of these natural structures. Throughout, the selfie camera and self-avatar features proved to be effective lubricants for student engagement.

**Figure 3.**
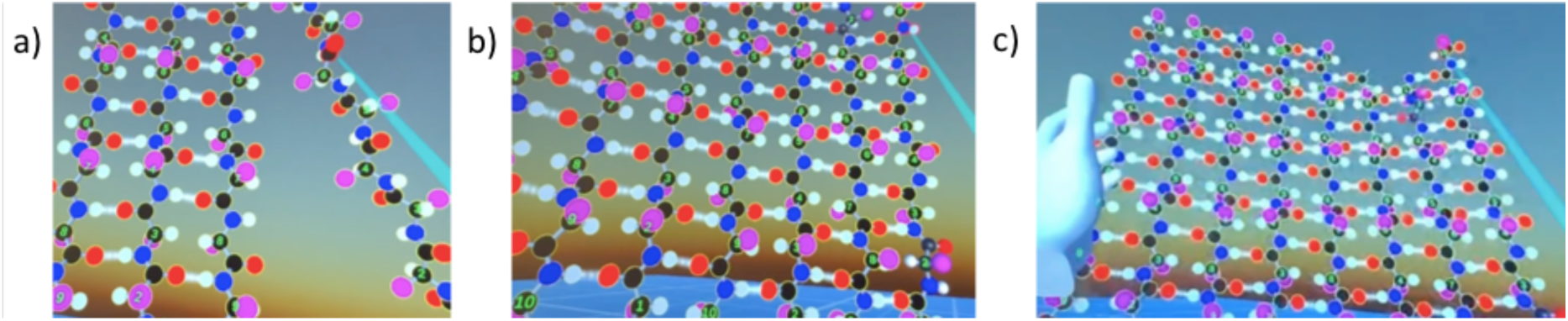
Example of a β-barrel built by a student. (a) The student has constructed an anti-parallel β-sheet using three decapeptides. A fourth is being brought in from the right. This quickly snaps into place from top to bottom with a zip-like smoothness, as the locked hydrogen bonds on the main sheet direct the conformation of the incoming peptide. (b) After four subsequent decapeptides have been added, an eight-peptide sheet is formed. It has taken less than ten minutes to build this structure. (c) Now the student is faced with trying to fold the entire structure in on itself so that a barrel can be formed. This proved to be too difficult – “*like trying to fold a bedsheet in the wind*”. However, the student learned a valuable lesson about both the strength and flexibility of the sheet. As well as stimulating discussion about other ways to complete the task, this experiment also gave us the impetus to incorporate the ‘freezing’ function into Peppy.

**Figure 4.**
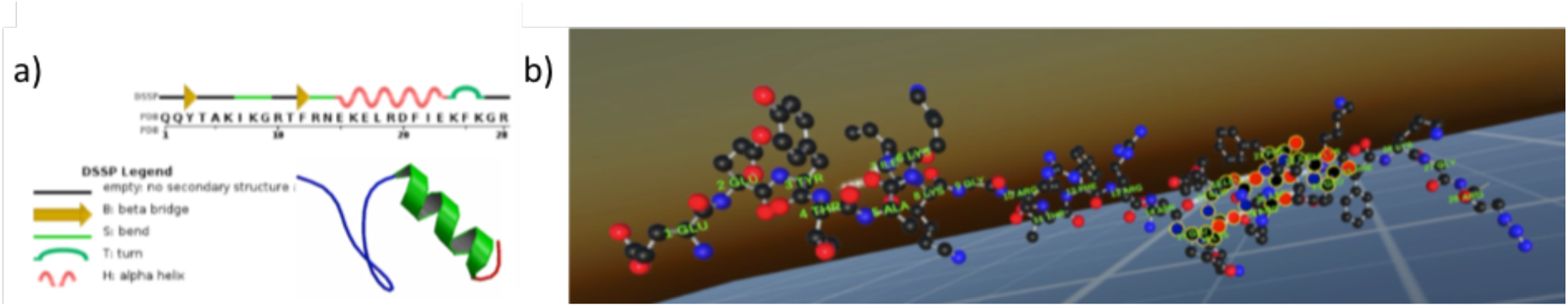
Example of a real peptide structure built by a student. (a) Sequence, secondary structure and cartoon structure of the 28-residue peptide, a section of zinc finger (PDB ID: 1FSV). After selecting residues 15-24 and adjusting the ϕ and ψ angle values on the Ramachandran plot to −57° and −47°, respectively, the student observed that section to smoothly settle into an α-helical configuration, with the hydrogen bonds (white) stabilising a rigid configuration. The student was then able to reflect on the orientation of the side-chains, the space that they occupy and the possible forces between them. Residues 1-14 did not so easily adopt a β-sheet, but this was valuable in helping the student think about what has to happen for proteins to fold. Indeed, had the ‘freeze’ function characteristic of the latest build of Peppy been available, the task of moving these residues to form their native configuration would have been simpler.

The students completed two assignments related to their experience with Peppy. The first was a standard laboratory report that provided proof that they had thoughtfully worked through the tasks. It took the form of screenshots embedded into a contextual narrative explaining the process, outcomes and lessons learned. Two examples of these are provided in the **Supplementary Material**.

The second task was more reflective and extrapolative. Students were asked to suggest workflows for the deployment of Peppy in 2^nd^ year classes, and were encouraged to propose features they would like to see added or changed. Many students more than fulfilled their obligations by trying to create ambitious and/or whimsical structures (**Figure 5**) or perform other manipulations which tested the limits of the software.

**Figure 5.**
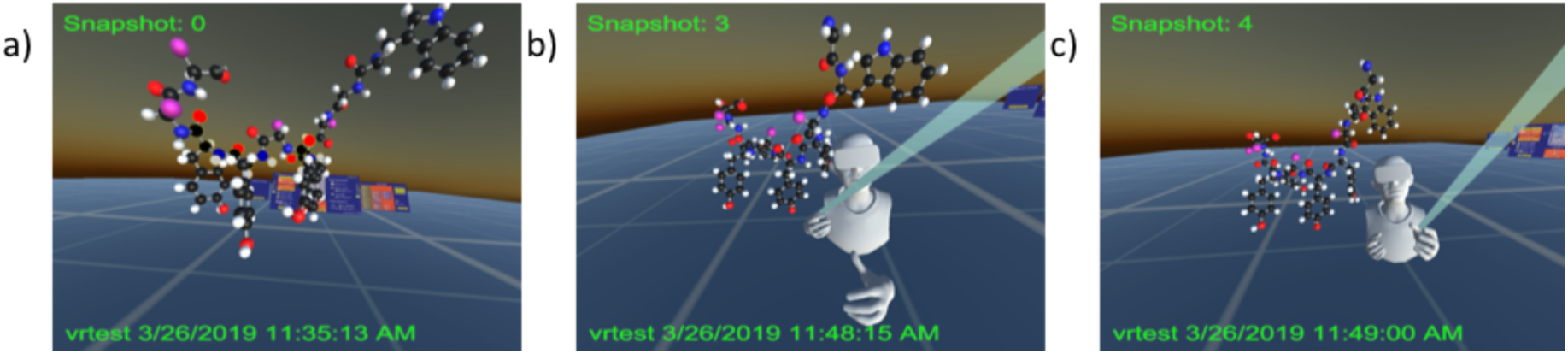
Examples of animals made by a student in the second task. (a) Initial attempt at a 10-residue peptide dog, which the student noted looks more like a giraffe due to the long neck. (b) Refined 10-residue peptide dog where the tryptophan head has been moved to residue 9 and residue 10 mutated to glycine to create an ear. (a) is a standard screen shot and(b) and (c) were taken using the selfie camera. The student noted that they liked image (c) because the dog appeared to be looking down upon their avatar. This exercise taught the student a lot about steric hinderance and the effect of ϕ and ψ angle values on overall structure.

During both testing phases, the students provided an abundant list of desirable features and noted problems with the existing implementation, many of which we added or fixed to arrive at the version reported herein, highlighting the agility of working within a game engine framework and the advantages of including front-line teaching academics in the development team.

**Figure 6.**
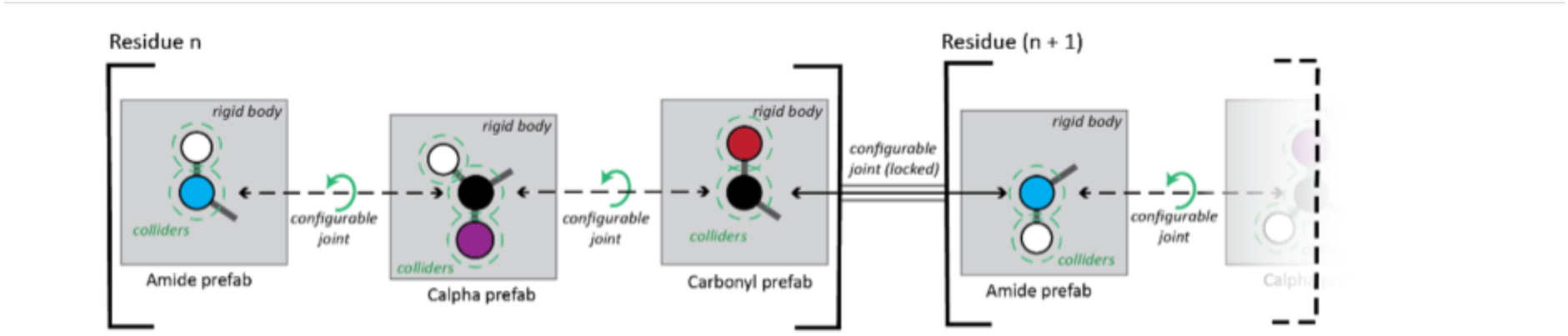
Illustration of the Peppy backbone architecture. The *n*^th^ amino acid comprises at least three rigidbody prefab units, representing the amide (N-H) and carbonyl (C=O) functional groups and the Calpha plus side chain (Cα + ‘R’) unit. The units are connected by configurable joints, but only those within a residue are freely rotatable (green arrows). Each atom is represented by a collider with a fixed radius specific to that atom type (green dashed lines), where type refers to the element as well as its chemical environment.

**Figure 7.**
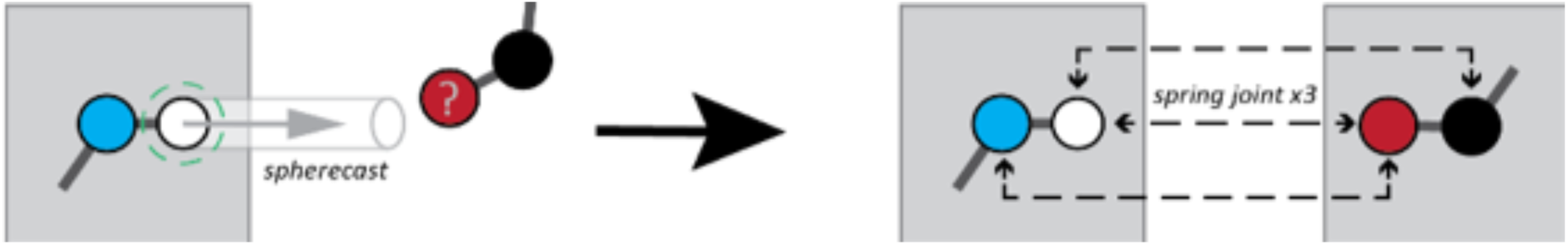
Illustration of hydrogen bond modelling in Peppy. Hydrogen bonding pairs are discovered by projecting in real time a cylinder from each donor group in line with the backbone N-H bond. If an acceptor is found, a hydrogen bond is modelled using three spring joints. The parameters of the cylinder and spring function are provided in **Supporting Information Table S6**.

**Figure 8.**
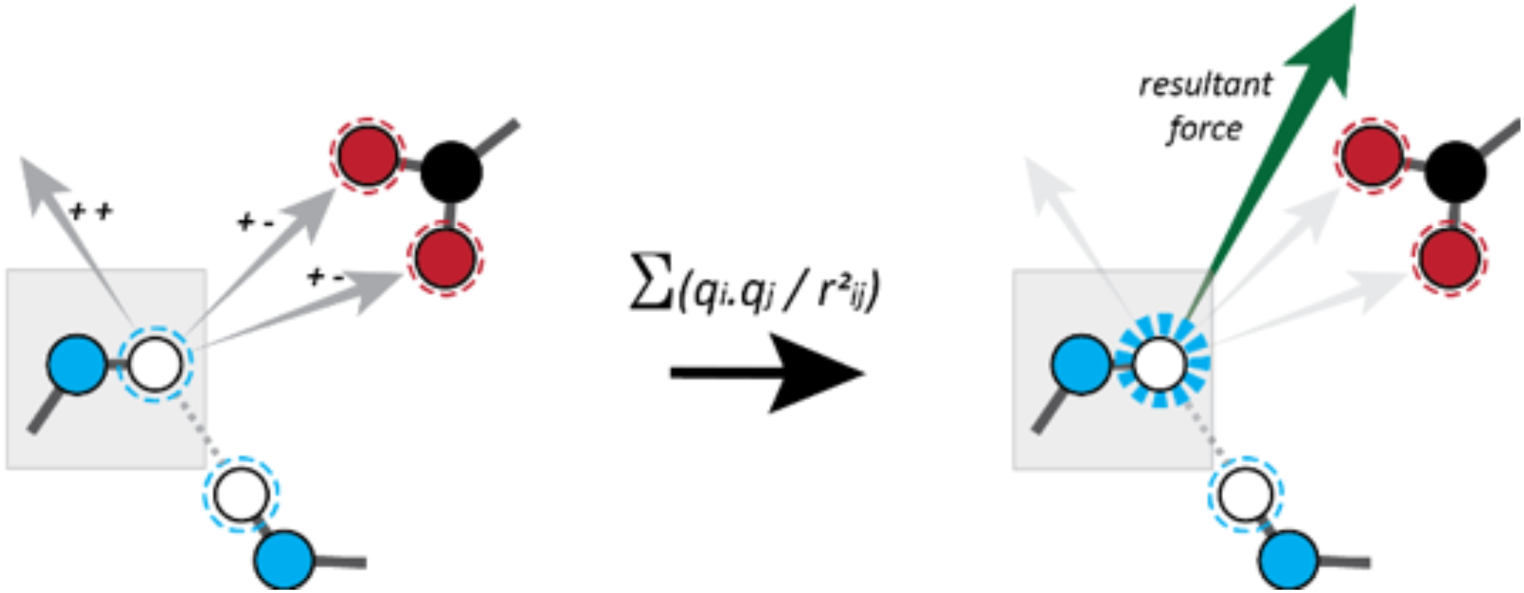
Illustration of electrostatic interaction modelling in Peppy. The total force on a partially charged atom is the sum of its Coulombic interactions with all other partially charged atoms (positively charged: white with blue dashed outline; negatively charged: red with red dashed outline).

## Conclusions

Our goal was to develop a VR application to facilitate teaching the principles of protein structure. We hypothesised that the experience of using VR would be progressive and would provide additional insight over what has previously been available, such as 2D printed images, 3D graphics projected onto 2D, and physical models. We harnessed the existing Unity video game physics engine and game development protocols to facilitate rapid prototyping and responsive development. We did not attempt to replicate the true underlying physical chemistry although we did take inspiration from existing modelling protocols such as molecular dynamics simulation force fields.

The resulting program, Peppy, presents the basic elements of protein secondary structure in an accessible format that is novel, dynamic and extremely tangible as well as fun to use. It is possible to easily and quickly investigate a wide range of conformational properties of the polypeptide backbone and sidechains. Remarkably, despite its simplicity, it is possible to build a large variety of complex multi-peptide structures using Peppy. In fact, the simplicity of the simulation becomes a strength when framed within the teaching and learning process as it provokes questions about the validity of the model. The physical parameters and assumptions of the underlying model are accessible and transparent and, as is so often the case when teaching with reference to a metaphor, higher levels of understanding and enlightenment result when there is appreciation of where the analogy is deficient. Indeed, students are specifically encouraged to test and probe the rules driving the simulation by changing the tuneable factors and exploring collisions and tensions. In all these ways, Peppy invites active challenge, questioning and critique by the students, all of which hopefully lead toward a deeper understanding of and interest in the forces and factors affecting protein structure.

## Supplementary Material

One file (Doak_ProtSci_SuppMat.pdf) containing two student reflections on using Peppy and eight supplementary tables of key parameter values.

## Acknowledgements

DGD, JAG and JRA acknowledge support from the University of Auckland Vice-Chancellor’s Strategic Development Fund. JAG and DGD also acknowledge support from the School of Chemical Sciences, University of Auckland. JRA is additionally supported by a Rutherford Discovery Fellowship (15-MAU-001). GSD is grateful to the following students for their permission to use examples of their work: Emily Thunberg, Michelle Meier, Zelda Weinstock and Andreas Orsmond.

## Notes

#### Summary of Updates

Minor formatting errors

## References

1. Balo AR, Wang M, Ernst OP (2017) Accessible virtual reality of biomolecular structural models using the Autodesk Molecule Viewer. Nat Methods 14:1122.

2. Bennie S, Ranaghan K, Deeks H, E. Goldsmith H, B. O’Connor M, J. Mulholland A, R. Glowacki D (2019) Teaching Enzyme Catalysis Using Interactive Molecular Dynamics in Virtual Reality. Journal of Chemical Education.

3. Bondi A (1964) van der Waals Volumes and Radii. The Journal of Physical Chemistry 68:441–451.

4. Borrel A, Fourches D (2017) RealityConvert: a tool for preparing 3D models of biochemical structures for augmented and virtual reality. Bioinformatics 33:3816–3818.

5. Goddard TD, Brilliant AA, Skillman TL, Vergenz S, Tyrwhitt-Drake J, Meng EC, Ferrin TE (2018) Molecular Visualization on the Holodeck. Journal of Molecular Biology 430:3982–3996.

6. Grebner C, Norrby M, Enström J, Nilsson I, Hogner A, Henriksson J, Westin J, Faramarzi F, Werner P, Boström J (2016) 3D-Lab: a collaborative web-based platform for molecular modeling. Future Medicinal Chemistry 8:1739–1752.

7. Humphrey W, Dalke A, Schulten K (1996) VMD: Visual molecular dynamics. J Mol Graph Model 14:33-38. PMID: WOS:A1996UH51500005 {Medline}

8. Johnston APR, Rae J, Ariotti N, Bailey B, Lilja A, Webb R, Ferguson C, Maher S, Davis TP, Webb RI, McGhee J, Parton RG (2018) Journey to the centre of the cell: Virtual reality immersion into scientific data. Traffic 19:105–110.

9. Kingsley LJ, Brunet V, Lelais G, McCloskey S, Milliken K, Leija E, Fuhs SR, Wang K, Zhou E, Spraggon G (2019) Development of a virtual reality platform for effective communication of structural data in drug discovery. Journal of Molecular Graphics and Modelling 89:234–241.

10. Kolasinski EM. Simulator Sickness in Virtual Environments. (1995).

11. Mikropoulos TA, Natsis A (2011) Educational virtual environments: A ten-year review of empirical research (1999–2009). Computers & Education 56:769–780.

12. Norrby M, Grebner C, Eriksson J, Boström J (2015) Molecular Rift: Virtual Reality for Drug Designers. J Chem Inf Model 55:2475–2484.

13. O’Connor M, Deeks HM, Dawn E, Metatla O, Roudaut A, Sutton M, Thomas LM, Glowacki BR, Sage R, Tew P, Wonnacott M, Bates P, Mulholland AJ, Glowacki DR (2018) Sampling molecular conformations and dynamics in a multiuser virtual reality framework. Science Advances 4:eaat2731.

14. O’Connor MB, Bennie SJ, Deeks HM, Jamieson-Binnie A, Jones AJ, Shannon RJ, Walters R, Mitchell TJ, Mulholland AJ, Glowacki DR (2019) Interactive molecular dynamics in virtual reality from quantum chemistry to drug binding: An open-source multi-person framework. The Journal of Chemical Physics 150:220901.

15. Ratamero EM, Bellini D, Dowson CG, Römer RA (2018) Touching proteins with virtual bare hands. J Comput Aid Mol Des 32:703–709.

16. Reif MM, Hünenberger PH, Oostenbrink C (2012) New Interaction Parameters for Charged Amino Acid Side Chains in the GROMOS Force Field. Journal of Chemical Theory and Computation 8:3705–3723.

17. Reif MM, Winger M, Oostenbrink C (2013) Testing of the GROMOS Force-Field Parameter Set 54A8: Structural Properties of Electrolyte Solutions, Lipid Bilayers, and Proteins. Journal of Chemical Theory and Computation 9:1247–1264. PMID: WOS:000315018300043 {Medline}

18. Salvadori A, Del Frate G, Pagliai M, Mancini G, Barone V (2016) Immersive virtual reality in computational chemistry: Applications to the analysis of QM and MM data. International Journal of Quantum Chemistry 116:1731–1746.

19. Schrodinger, LLC. The PyMOL Molecular Graphics System, Version 1.8. (2015).

20. Shahbazi Z (2015) Mechanical Model of Hydrogen Bonds in Protein Molecules. American Journal of Mechanical Engineering 3:47–54.

21. Stefani C, Lacy-Hulbert A, Skillman T (2018) ConfocalVR: Immersive Visualization for Confocal Microscopy. Journal of Molecular Biology 430:4028–4035.

22. Stone JE, Kohlmeyer A, Vandivort KL, Schulten K. Immersive Molecular Visualization and Interactive Modeling with Commodity Hardware. In: Bebis G, Boyle R, Parvin B, Koracin D, Chung R, Hammound R, Hussain M, Kar-Han T, Crawfis R, Thalmann D, Kao D, Avila L, Eds. (2010) Advances in Visual Computing. Springer Berlin Heidelberg, Berlin, Heidelberg, pp. 382–393.

23. Trindade J, Fiolhais C, Almeida L (2002) Science learning in virtual environments: a descriptive study. British Journal of Educational Technology 33:471–488.

24. Zheng M, Waller MP (2017) ChemPreview: an augmented reality-based molecular interface. Journal of Molecular Graphics and Modelling 73:18–23.

