## Supplementary Information for "Peppy: A Virtual Reality Environment for Exploring the Principles of Polypeptide Structure"

**Table S1. Atomic radii and masses used in Peppy.** The base radii of the colliders for different elements are listed. There is also a global scaling factor which is used to scale all colliders, which has a default value of 0.675. The atomic radii were taken from Bondi's compilation(1) and the units changed from Ångstroms to metres to reflect the human scale of the molecules in Peppy. Similarly, the masses are in kilograms.

| Element | Radius (m) | Mass (kg) |
|---|---|---|
| C | 0.170 | 1.2 |
| H | 0.120 | 0.1 |
| N | 0.155 | 1.4 |
| O | 0.152 | 1.6 |
| S | 0.180 | 3.2 |
| R | 0.180[a] | 1.5[a] |

[a] The Cα-R radius and mass are arbitrary since R is a dummy atom used to represent any side chain.

**Table S2. Bond lengths and angles within each rigidbody prefab unit.** These bonds and angles are fixed and thus there are not associated force constants. While the actual bond length units are Ångstroms, metres are appropriate given the human scale of the molecules in Peppy.

| Prefab unit | Bond type | Bond length (m) | Angle type | Angle (degrees) |
|---|---|---|---|---|
| Amide | N-H | 0.100 | | |
| Calpha | Cα-H | 0.100 | H-Cα-R | 110 |
| | Cα-R | 0.150[a] | | |
| Carbonyl | C=O | 0.124 | | |

[a] The Cα-R bond length is arbitrary since R is a dummy atom used to represent any side chain.

**Table S3. Bond length, target dihedral angle and torque values for the peptide backbone configurable joints that connect the prefab units.** The target dihedral angle and associated torque values for the ϕ (Cα-N) and ψ (Cα-C) dihedral angles can be tuned by the user within the specified range.

| Bond type | Bond length (m) | Target dihedral angle (degrees) | Torque (N·m) |
|---|---|---|---|
| Peptide (C-N) | 0.133 | fixed [0] | n/a |
| Amide-Calpha (Cα-N) | 0.146 | range [-180, +180] | range [0 – 400] |
| Calpha-Carbonyl (Cα-C) | 0.151 | range [-180, +180] | range [0 – 400] |

**Table S4. Bond angles for the peptide backbone configurable joints that connect the prefab units.**

| Angle type | Angle value (degrees) |
|---|---|
| Cα-C-O | 120.5 |
| Cα-C-N | 116.0 |
| O-C-N | 123.5 |
| C-N-H | 119.5 |
| C-N-Cα | 122.0 |
| H-N-Cα | 118.5 |
| N-Cα-C | 111.0 |

**Table S5. Sidechain Geometry.** For simplicity, most bond lengths between sidechain heavy atoms are equivalent. Only bond angle departures from idealised $sp^2$ (trigonal) geometry are listed explicitly.

| Amino acid | Bond type | Bond length (m) |
|---|---|---|
| All | X-X [a] | 0.150 |
| | X-H | 0.100 |
| Aspartate, Glutamate | $C\text{-}O^-$ | 0.125 |
| Asparagine, Glutamine | C=O | 0.125 |

| Amino acid | Angle type | | | Angle value (degrees) |
|---|---|---|---|---|
| Tryptophan | CB | CG | CD1 | 126.0 |
| | CG | CD1 | NE1 | 107.0 |
| | CD1 | NE1 | CE2 | 110.0 |
| | NE1 | CE2 | CD2 | 108.0 |
| | CE2 | CD2 | CG | 108.0 |
| | CD2 | CG | CD1 | 107.0 |
| | CG | CD2 | CE3 | 132.0 |
| | NE1 | CE2 | CZ2 | 132.0 |
| Histidine | CB | CG | CD2 | 126.0 |
| | CG | CD2 | NE2 | 107.0 |
| | CD2 | NE2 | CE1 | 107.0 |
| | NE2 | CE1 | ND1 | 108.0 |
| | CE1 | ND1 | CG | 110.0 |

[a] Atom type X is C, N, O or S.

**Table S6. Desired, minimum and maximum interatomic distances, spring force constants and damping constants for the dynamic hydrogen bond configurable joints.** 'Inner' refers to the hydrogen – acceptor distance, and 'outer' to the distance between the donor and the atom directly bonded to the acceptor.

| **Hydrogen bond type** | $r_{min}$ **(m)** | $r_{max}$ **(m)** | $k_{hb}$ ***(N/m)*** | $d_{hb}$ ***(rad/s)*** |
|---|---|---|---|---|
| Inner | *–infinity* | 0.16 | range [0 – 2500] | 5 |
| Outer | 0.35 | *infinity* | range [0 – 2500] | 5 |

**Table S7. Atomic partial charges for calculating electrostatic interactions.** Atom type codes and associated partial charges are simplified versions of those used in the GROMOS 54A8 force field(2; 3).

| **Amino acid unit** | **Atom type** | ***Q (e = -1)*** |
|---|---|---|
| Backbone | H | +0.25 |
| | O | -0.25 |
| Aspartate | OD1 | -0.50 |
| | OD2 | -0.50 |
| Glutamate | OE1 | -0.50 |
| | OE2 | -0.50 |
| Arginine | CZ | +0.34 |
| | NH1 | +0.33 |
| | NH2 | +0.33 |
| Lysine | NZ | +0.25 |
| | HZ1 | +0.25 |
| | HZ2 | +0.25 |
| | HZ3 | +0.25 |
| Histidine | ND1 | +0.50 |
| | NE2 | +0.50 |

**Table S8. Drag factor values for rigidbody dynamics.**

| Drag slider/user value | Translational drag factor | Angular drag factor |
|---|---|---|
| 0 | 5 | 5 |
| 100 | 25 | 25 |
| "Frozen" | infinity | infinity |

### References


1. Bondi A (1964) van der Waals Volumes and Radii. The Journal of Physical Chemistry 68:441-451.
2. Reif MM, Hünenberger PH, Oostenbrink C (2012) New Interaction Parameters for Charged Amino Acid Side Chains in the GROMOS Force Field. Journal of Chemical Theory and Computation 8:3705-3723.
3. Reif MM, Winger M, Oostenbrink C (2013) Testing of the GROMOS Force-Field Parameter Set 54A8: Structural Properties of Electrolyte Solutions, Lipid Bilayers, and Proteins. Journal of Chemical Theory and Computation 9:1247-1264. PMID: WOS:000315018300043 {Medline}
